# Predicting the cross-continental spread of the cassava brown streak disease epidemic in sub-Saharan Africa

**DOI:** 10.1101/2025.10.10.681618

**Authors:** David Godding, Richard O. J. H. Stutt, Merveille Koissi Savi, Corneille Ahanhanzo, Fidèle Tiendrebeogo, Oumar Doungous, Monde Godefroid, Zeyimo Bakelana, Jacques François Mavoungou, Allen Oppong, Elijah Ateka, Willard Mbewe, Nurbibi Cossa, Chukwuemeka Nkere, Ibrahim Umar Mohammed, Marie Claire Kanyange, Alusaine Edward Samura, Adjata Djodji, Titus Alicai, Patrick Chiza Chikoti, Fred Tairo, Peter Sseruwagi, Joseph Ndunguru, Angela Obiageli Eni, Justin Simon Pita, Christopher A. Gilligan

## Abstract

Cassava brown streak disease (CBSD) is a major threat to smallholder farmers in sub-Saharan Africa (SSA), where cassava is a staple crop. Caused by cassava brown streak ipomoviruses (CBSIs), CBSD has spread extensively since its detection in Uganda in 2004, raising concerns about ongoing spread through Southern and Central Africa and potential expansion to West Africa, home to the world’s largest cassava producer, Nigeria.

Building on a stochastic epidemiological model that predicts CBSD spread at the scale of Uganda, we incorporate extensive field surveillance records to extend the model to all thirty-two major cassava-producing countries in SSA. We then deploy the model to address key strategic questions such as estimating the present day CBSD distribution and predicting rates of ongoing spread towards West Africa. We also evaluate the risk of direct introductions resulting from long range movement of infected planting material, which could trigger outbreaks beyond predicted localised dispersal limits. Our model predicts the likely arrival of CBSD in Nigeria via cross-continental spread within 25 years, and if directly introduced anywhere in West Africa, spreading to most West African nations within 10 years.

The risks of ongoing and future CBSD spread highlighted in this study underscore the need for proactive phytosanitation measures, including clean seed programs, vector control, and quarantine policies to curb CBSD spread. Moreover, the model described in this study not only provides estimates for arrival times across SSA, but also lays the foundations for a continental-scale quantitative framework wherein both surveillance and management options can be explored and optimised.

## 1 Introduction

Emerging epidemics and pest infestations of agricultural crops are a major burden for smallholder farmers in sub-Saharan Africa (SSA), causing income loss and exacerbating food security (Harvey *et al*., 2014). Smallholder farmers often have limited access to resistant varieties and pesticides to mitigate the continually shifting burden of pests and pathogens, in contrast to industrialised agriculture (Gomez y Paloma *et al*., 2020). Therefore, smallholder farmers are most vulnerable to the yield losses that novel pests and pathogens can cause. Current and recent threats to staple crops in SSA include a range of transboundary pests and pathogens that spread within and between countries. Some, such as desert locusts (FAO, 2021) experience population explosions under favourable environmental conditions, leading to large swarms that disperse over long distances (Meynard *et al*., 2017; Retkute *et al*., 2021). Fall armyworm also spread rapidly across national boundaries (Day *et al*., 2017) following the first introductions in West and Central Africa (Goergen *et al*., 2016). Other disease-causing pathogens, such as cassava mosaic virus and cassava brown streak virus are less explosive but capable of spreading over long distances by human-mediated and insect vector transmission (Legg *et al*., 2011, 2015b, a,b; Alicai *et al*., 2019; Godding *et al*., 2023).

Slower spreading outbreaks allow time to prepare surveillance, early-warning, and mitigation strategies in disease-free regions prior to invasion. Predicting likely arrival times and spread within a new region is an essential component for forward planning. Here we focus on cassava brown streak disease (CBSD) and investigate the use of stochastic, epidemiological modelling to assess the risk to West and Central Africa from cross continental spread and direct introduction.

Since its first detection in Uganda in 2004, an epidemic of cassava brown streak disease has spread over 1000 km from Uganda (Alicai *et al*., 2007, 2019) to Rwanda (Munganyinka *et al*., 2018), Burundi (Alicai *et al*., 2007), western Kenya (Mware *et al*., 2009), the lake-zone of Tanzania (Mbanzibwa *et al*., 2011), central Democratic Republic of Congo (Muhindo *et al*., 2020a), and Zambia (Mulenga *et al*., 2018). The CBSD epidemic is having a major impact on the consumable and marketable yields of cassava for smallholder farmers, which threatens food security and economic development (Legg *et al*., 2015b) and is continuing to spread, threatening production in West Africa, which includes the world’s largest producer, Nigeria (FAO, 2022).

The CBSD epidemic is caused by single-stranded ipomoviruses, Cassava brown streak virus (CBSV) and Ugandan cassava brown streak virus (UCBSV), of the family *Potyviridae* (Tomlinson, K. *et al*., 2017), henceforth referred to collectively as cassava brown streak ipomoviruses (CBSIs). Cassava is vegetatively propagated, with stem cuttings taken from plants in one season used as planting materials in the next season, rather than planting from seed. As a result, in the absence of management efforts, once cassava plants in a field becomes infected with CBSD, they remain infected. Currently, there are no cassava varieties that are fully resistant to CBSD in SSA (Tumwegamire *et al*., 2018), and all West African varieties tested to date are susceptible to infection (Elegba *et al*., 2020; Ano *et al*., 2021). However, there are significant efforts to identify sources of resistance that are showing promise (Sheat *et al*., 2019). Until resistant varieties are widely available, management options at the larger scale involve phytosanitation programmes. These include clean seed programmes to disseminate uninfected cuttings and quarantine systems to reduce the movement of infected planting material (Legg *et al*., 2015b). At the field level, options include: vector control with pesticides (Omongo *et al*., 2022) to reduce transmission rates; roguing to remove infected plants; and selection of asymptomatic cuttings for propagation to bias the choice of cuttings away from those that are infected (Legg *et al*., 2015b). Each of these management options has different requirements in terms of investment and training, and varying probabilities of success according to the epidemiological and social context.

Cassava brown streak virus is spread by an insect vector, *Bemisia tabaci,* and by human-mediated movement of infected cuttings. Vector activity operates primarily at a local scale to spread infection, with cutting movement introducing the pathogen to new areas. The majority of cuttings are sourced from farmers’ own fields, but it is not uncommon to source from within a 10 km radius from friends and relatives (Szyniszewska *et al*., 2019). More rarely, farmers may source materials from further afield (i.e.,>50 km), which are likely from either traders or not-for-profit organisations (e.g., NGOs) (Szyniszewska *et al*., 2019). Therefore, when assessing how CBSD can spread to new regions, two distinct scenarios emerge. The first scenario captures the cumulative effect of relatively local and continuous spread with a strong local bias and occasional medium-range dispersal up to ∼50 km, resulting from the combined effect of vector activity and local cutting movements (Szyniszewska, 2020; Godding *et al*., 2023). The second scenario accounts for the risk of direct introduction of infected cuttings to a previously disease-free region over an unbounded distance (e.g., air, sea, or long-range land transport of cuttings), which could result in the direct introduction of CBSI to West Africa at any time. Moreover, in addition to the risk of direct introduction by private actors via sea or airports, it is not unprecedented for NGOs to move cassava cuttings by air (ICRC, 2011). Whilst such direct introduction events are likely to be rare due to the relative infrequency of attempted long-range movements, a single event could cause a large-scale independent outbreak.

The sparse reports of CBSD spread through Eastern, Southern and Central Africa do not give a full picture of the true current scale of the CBSD epidemic in SSA, especially of the risks to countries in Central and West Africa, where cassava is a major staple crop for both rural and urban populations. While CBSIs have been recorded in the Democratic Republic of Congo (DRC) (Muhindo *et al*., 2020a) and Zambia (Mulenga *et al*., 2018), to date, there are no epidemiological modelling studies to predict the future spread of CBSIs to guide policy makers across Central and West Africa in preparedness, monitoring, and management efforts. Godding et al. (Godding *et al*., 2023) recently analysed the spread of CBSD in Uganda in order to estimate dispersal parameters at country-wide scales. The stochastic, spatiotemporal model takes into account estimates of the density and connectivity of the cassava crop and the whitefly vector in influencing pathogen spread. The model, which was resolved to 1 km^2^, successfully predicted rates and extents of spread of CBSD throughout Uganda during the period 2004-2017 (Godding *et al*., 2023), for which well-documented surveillance data were available (Alicai *et al*., 2019). In this paper, we extend and evaluate the model developed in Uganda to predict likely cross-continental spread rates for all thirty-two major cassava-producing countries of SSA. Specifically, we use the spatially extended model to address the following questions:

- How accurately does the model predict the post-2004 regional spread of CBSD?
- How can additional information in the form of real-world reports of CBSD spread be incorporated into the model to reduce the uncertainty of predictions?
- What would be the impact of the direct introduction of CBSI-infected planting material to disease-free regions of West Africa?
- What is the current spatial distribution of the CBSD epidemic?
- What are the predicted arrival times for CBSIs to currently disease-free countries across sub-Saharan Africa in the absence of long-range direct introduction to disease-free regions?
- How much uncertainty is there in predictions of cross-continental spread?

In answering some of these questions, we also introduce innovations to reduce uncertainty in predicting risk by selecting epidemiological trajectories from large ensembles of simulations that account for limited positive records of disease occurrence in a sparsely sampled heterogeneous landscape.

## 2 Results

### 2.1 Validating predictions of regional spread from 2004 to the present day

Since the first observations of CBSD in Uganda in 2004, the epidemic has rapidly spread within and beyond Uganda. Previous studies have reported the strong correspondence of the model to surveillance data within Uganda during model validation (Godding *et al*., 2023). Beyond Uganda, a limited number of reports document CBSD arrival in previously disease-free regions (Alicai *et al*., 2007; Mulimbi *et al*., 2012; Mulenga *et al*., 2018; Munganyinka *et al*., 2018; Alicai *et al*., 2019; Muhindo *et al*., 2020a; Casinga *et al*., 2020) (Figure 1a).

**Figure 1:**
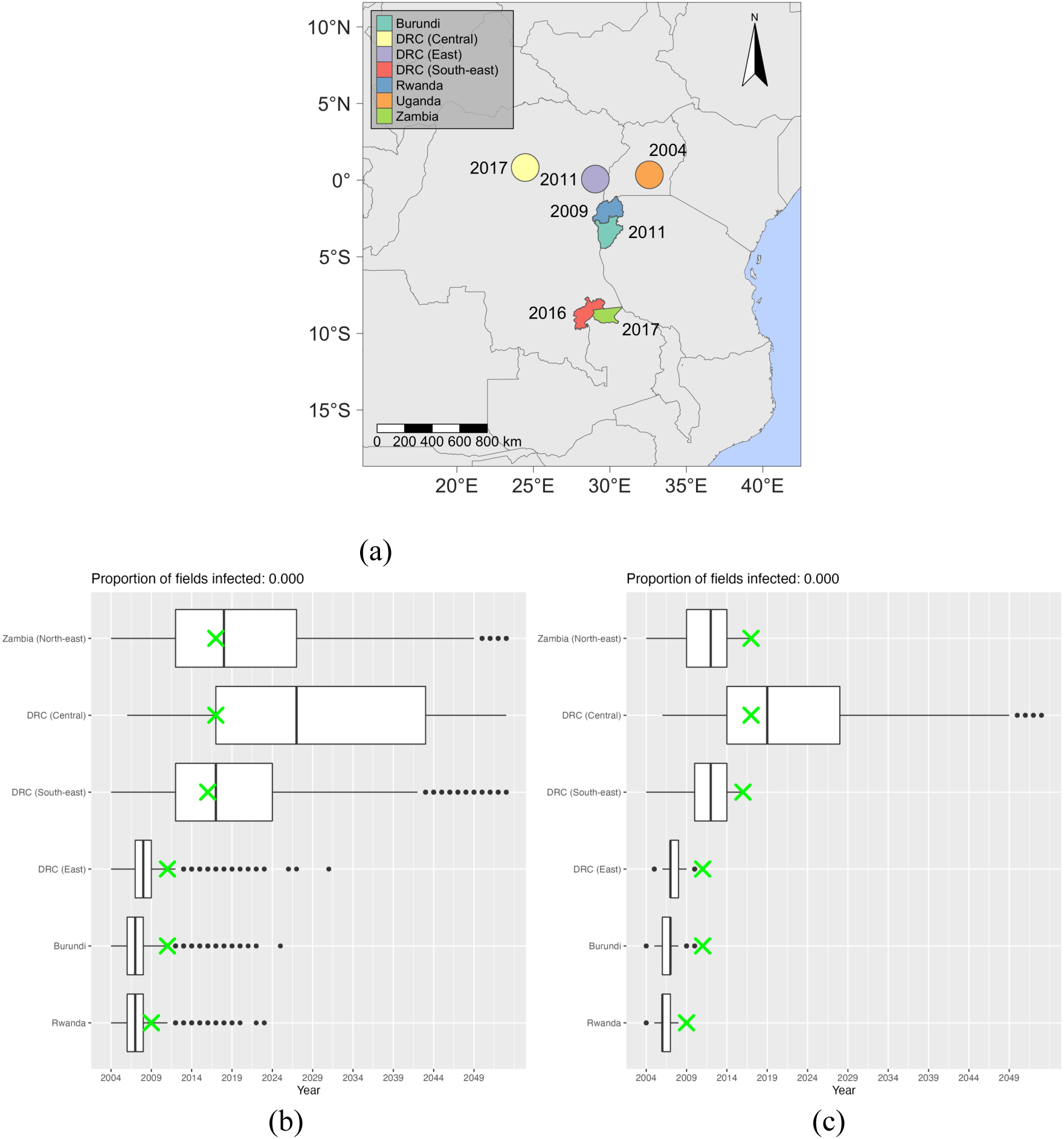
(a): Key observations of CBSD spread since the initial outbreak in Uganda in 2004. (b): Predicted arrival times in target regions compared with timing of real-world reports generated from all 9072 simulations initialised with infection only present at the sites of the first finding of CBSD in Uganda in 2004. Boxes indicate the median and the 25th and 75th percentiles; green crosses indicate year of real-world reporting. (c): Predicted arrival times in target regions compared with timing of real-world reports generated from a subset of 991 simulations out of the full set of 9072 simulations that corresponded to CBSD spread in five of the six regions (all except DRC (Central)) and surveillance data within Uganda.

From an ensemble of 9072 stochastic simulations for the spread of CBSI, beginning in 2004 within Uganda, we contrast the predicted epidemic arrival times with the real-world reports of CBSD detection (Figure 1b). As the underlying model (Godding *et al*., 2023) we would expect the model to predict CBSD arrival prior to or in the same year as the real-world reporting, with the duration between arrival of the pathogen and reports of disease presence being inversely related to the intensity of surveillance efforts. For five of the six CBSD regions, the model predictions correspond closely with the real-world reporting year. In the outlier case of Central DRC in which the report is comparatively early, the report still lies within the 25th percentile of simulations. Notably, the southeast DRC and Zambia reports in 2016 and 2017, respectively, are both approximately 1000 km from the original reports of the 2004 Ugandan outbreak. Nonetheless, the model predicts median arrival time within 1 year of the real-world report.

For forward predictions we then incorporate additional information on real-world spread on arrival time predictions. We generate a present-day infection status for all cassava hosts (including those unsurveyed) to act as the initial conditions for. In order to generate the full present day infection status, we take simulations initialised from the 2004 status and run forward to the present day, and then select those simulations consistent with surveyed infection status to act as the present-day initial conditions for forward simulation (see Methods and Discussion for further information). Specifically, we restrict the full set of unrestricted simulations to a subset of 991 simulations that correspond to verified reports of the disease within Uganda, Rwanda, Burundi, east DRC, south DRC and Zambia, but we do not constrain on the central DRC report. Contrasting Figure 1b with Figure 1c, we see the improved prediction accuracy for arrival times in DRC (central) (reducing lag of the median simulation from 10 to 2 years) because of incorporating additional information on real-world observations.

### 2.2 Direct introduction of CBSI to disease-free regions

In order to explore the impact of CBSI being introduced to different regions in West Africa in the absence of management efforts, we performed a simulated introduction of CBSI infected material at five independently selected cassava fields close to each of three major and well-connected cities in West Africa: Lagos, Nigeria; Abidjan, Côte d’Ivoire; and Douala, Cameroon (see Methods for definition of field and details of infection seeding methodology).

In the Nigeria scenario, simulations show rapid spread across southern Nigeria and into Benin and Ghana by year 3, with over 10% of cassava production affected, reflecting high cassava densities in these countries, with lower levels of infection in Togo and Cameroon where cassava density and connectivity is lower (Figure 2a). After 10 years, the epidemic is predicted to have bulked up rapidly in the three countries (Nigeria, Benin and Ghana) that experienced rapid spread within 3 years but not greatly expanded beyond this region (Figure 2b).

**Figure 2:**
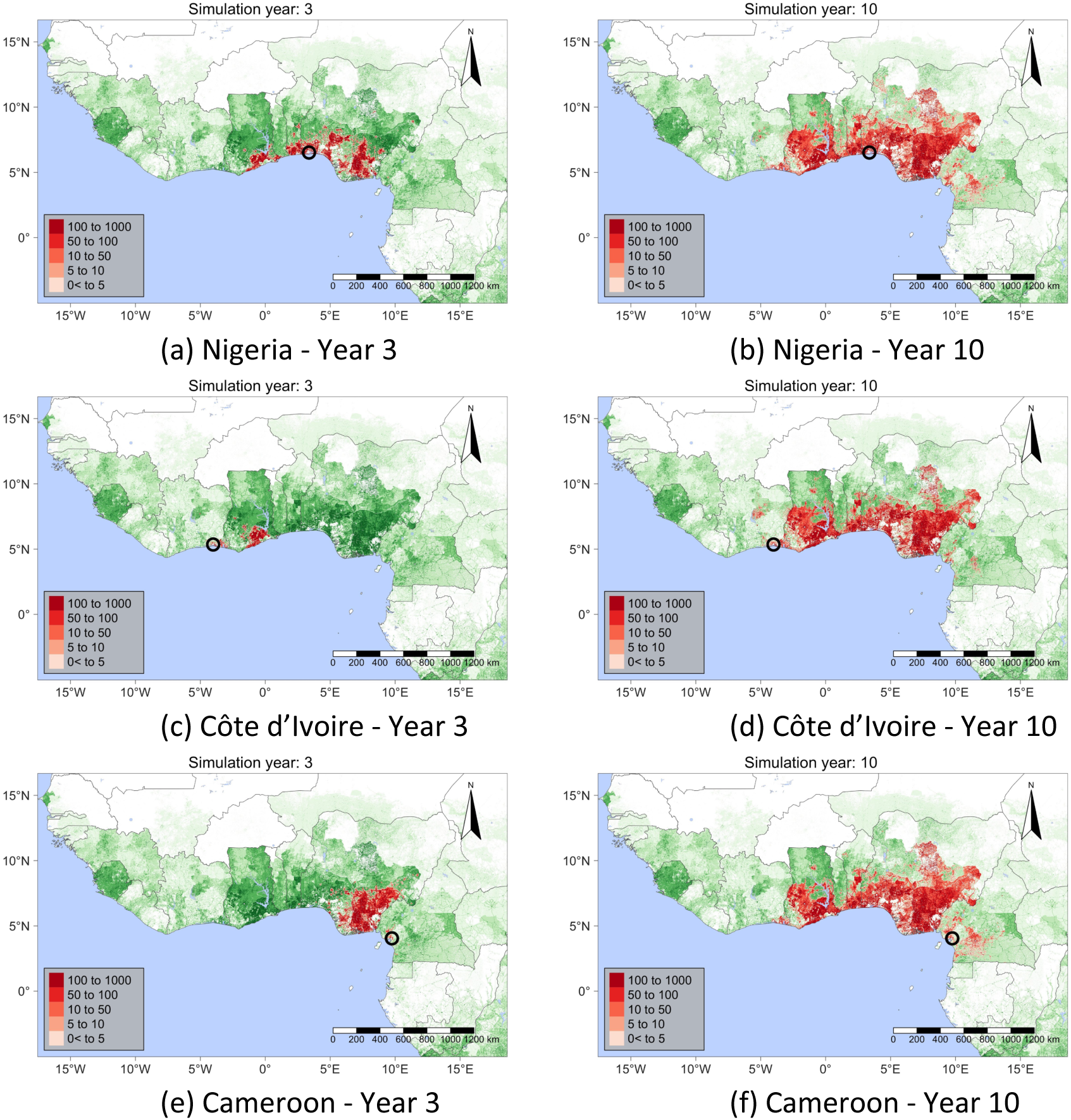
A single realisation of the model showing the median simulation of epidemic spread resulting from the direct introduction of infected cuttings (equivalent of five separate cassava fields – see Methods) to the area surrounding major and well-connected cities in West Africa. Black circles indicate the area in which initial infected fields were located.

**Figure 3:**
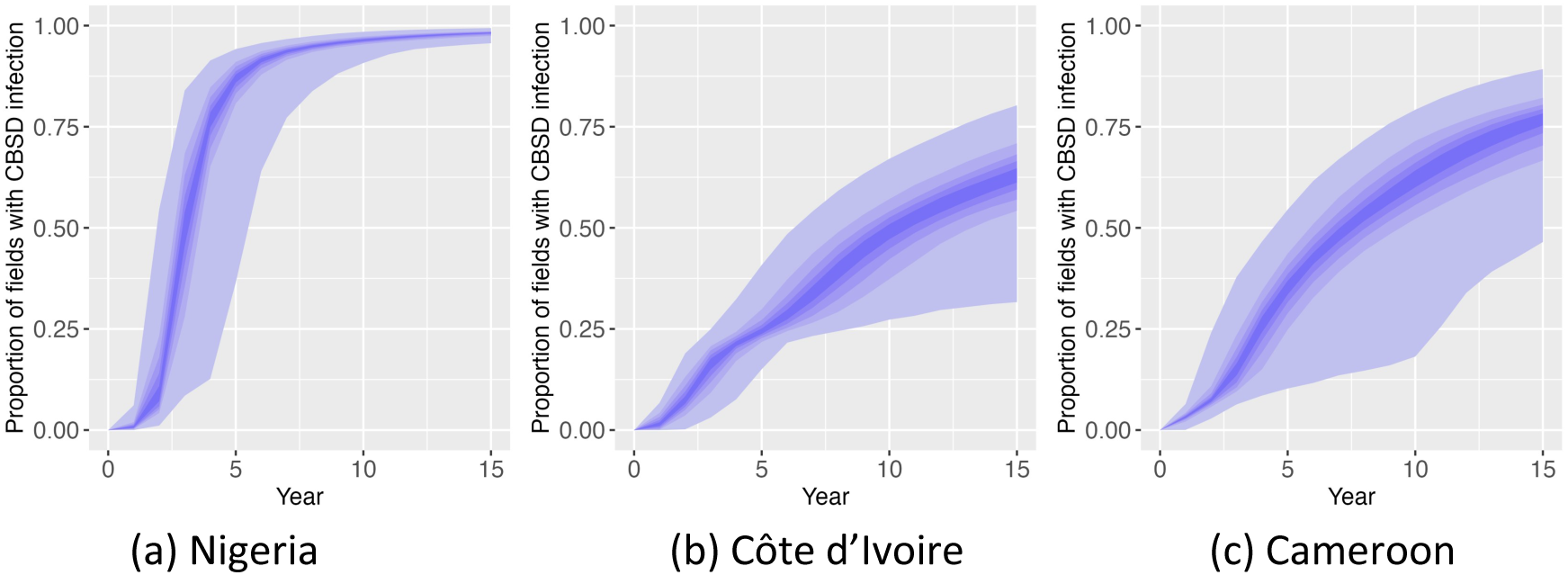
Disease progress curves (DPCs) from approximately 1000 direct introduction simulations wherein the equivalent of five separate infected cassava fields were introduced to the area surrounding major and well-connected cities in West Africa. DPCs restrict calculations of disease bulk up to the respective country of disease introduction. Each gradation in shading indicates ten percentiles of the distribution of infection levels, with the exception of the central band encompassing the 40-60 percentile range.

With epidemic initialisation in Côte d’Ivoire, we see relatively slow bulk up within Côte d’Ivoire by year 3 but the epidemic rapidly spreads in southern Ghana (Figure 2c). By year 10, the model predictions indicate that the epidemic has spread extensively into countries to the east and the distribution appears almost indistinguishable from the scenario where infection is initialised in Nigeria (Figure 2d). In the Cameroon scenario, infection bulks up relatively slowly within country, but rapidly spreads in southeastern Nigeria (Figure 2e), with the similar regional distribution emerging by year 10 as observed in Nigeria and Côte d’Ivoire scenarios (Figure 2f). The initial site of direct introduction appears to have little impact on the 10-year spatial distribution of infected fields due to the high density and connectivity of cassava cropping across this region of West Africa (Figures 2b, d, f and 3), with all introduction sites reaching at least 50% prevalence within this timeframe.

### 2.3 Predicting the present-day distribution and future spread of the epidemic

Given the extreme sparsity of CBSD surveillance data relative to the scale of the epidemic, the exact present-day distribution of the epidemic is not known. From a set of 9072 simulations covering the period from 2004 to 2054, we isolate the subset of 414 simulations that correspond to all real-world reports of CBSD spread (Figure 1a) and bulk up within Uganda (Alicai *et al*., 2007; Mulimbi *et al*., 2012; Mulenga *et al*., 2018; Munganyinka *et al*., 2018; Alicai *et al*., 2019; Muhindo *et al*., 2020a; Casinga *et al*., 2020).. Each simulation initialised in 2004 and consistent with survey results to the present day forms a potential complete infection status (critically, for unsurveyed locations) for the whole continent at the present day (see Methods for further information). The predicted present day (taken as 2025) CBSD distribution incorporates the historically endemic region of coastal East Africa and Malawi as well as the newly endemic regions of Uganda, western Kenya, north-western Tanzania, Rwanda, Burundi, and Zambia (Figure 4a). Considering the post-2004 spread west of Uganda up to the present day, our simulation results indicate that the majority of eastern DRC is highly likely (probability above 50%) to be endemic, with a high probability of presence in the region surrounding Kisangani in north-central DRC. Most notably, predictions indicate likely spread further west of current reports, specifically extensive spread across southern DRC (probability above 40%) and into north-eastern Angola (probability above 20%).

**Figure 4:**
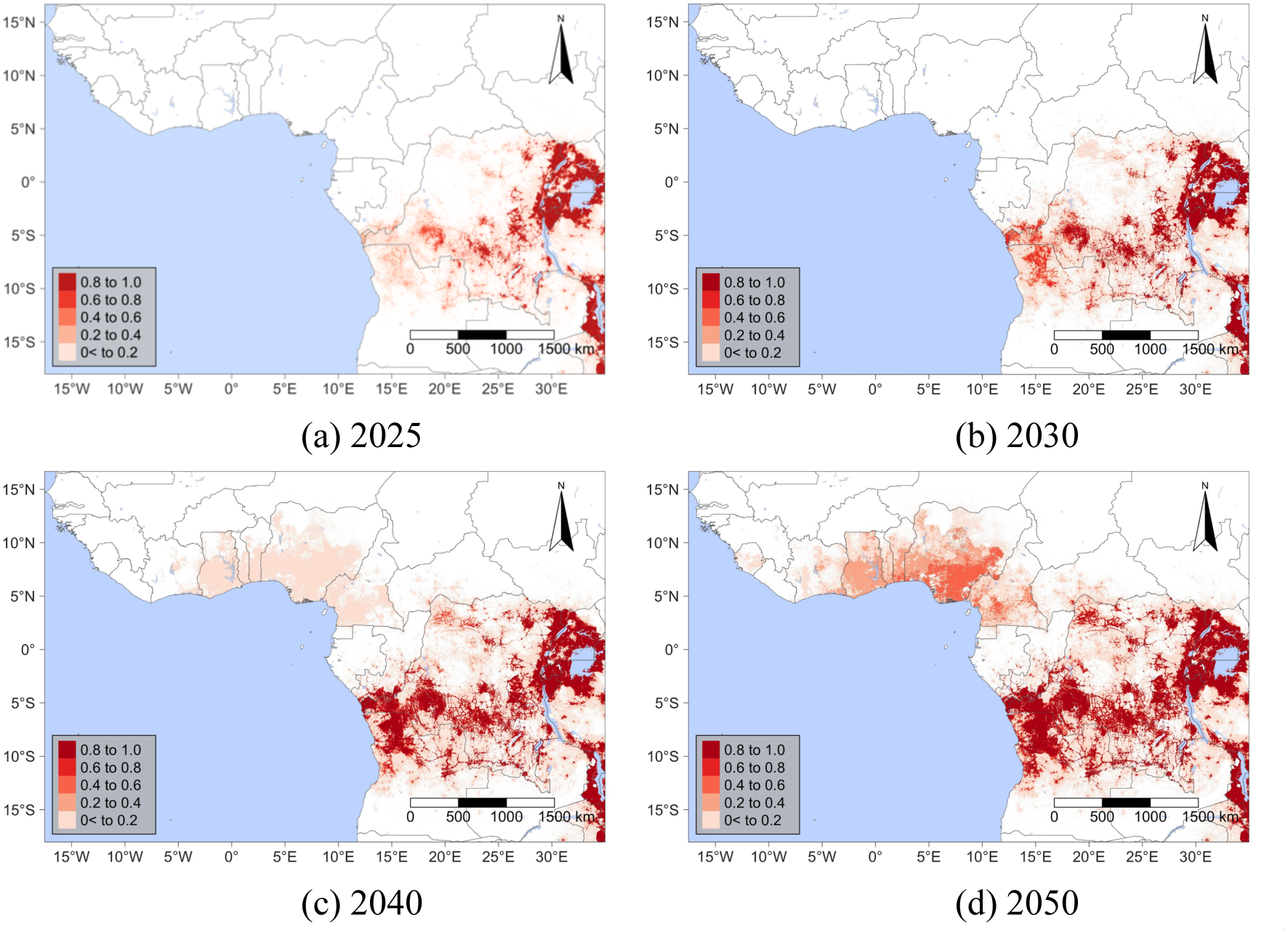
Risk maps representing the probability that any fields within a given one km^2^ raster cell will be infected with CBSD from (a): the present day (2025) through subsequent decades to (d): 2050.

We now consider the predicted 2030 epidemic distribution, with risk maps highlighting extensive epidemic expansion beyond the present day predicted distribution in northern Angola, south-western DRC, and north-central DRC, with more than a fivefold increase in expected area infected from the 2025 prediction. Likely new regions include Cabinda in Angola and southern Congo-Brazzaville (Figure 4b). Figure 5a highlights the major regions of change predicted by the simulations in the form of a differential risk map, which is derived by subtracting the risk map in 2025 from the risk map in 2030. By 2040, risk maps show expansion in north-western DRC and new incursions in the Central African Republic (CAR) and Gabon. Stochastic model simulations indicate approximately a 15% chance of CBSI having arrived in Cameroon and Nigeria, with potential spread as far west as Côte d’Ivoire (Figure 4c and 5b). By 2050, in the absence of effective control programmes, the model predicts extensive epidemic spread in West Africa, with 45% of simulations reaching Nigeria (Figure 4d and 5c). In addition, Supplementary Movie 1 illustrates a representative single realisation of the spatiotemporal dynamics of CBSD spread across the continent.

**Figure 5:**
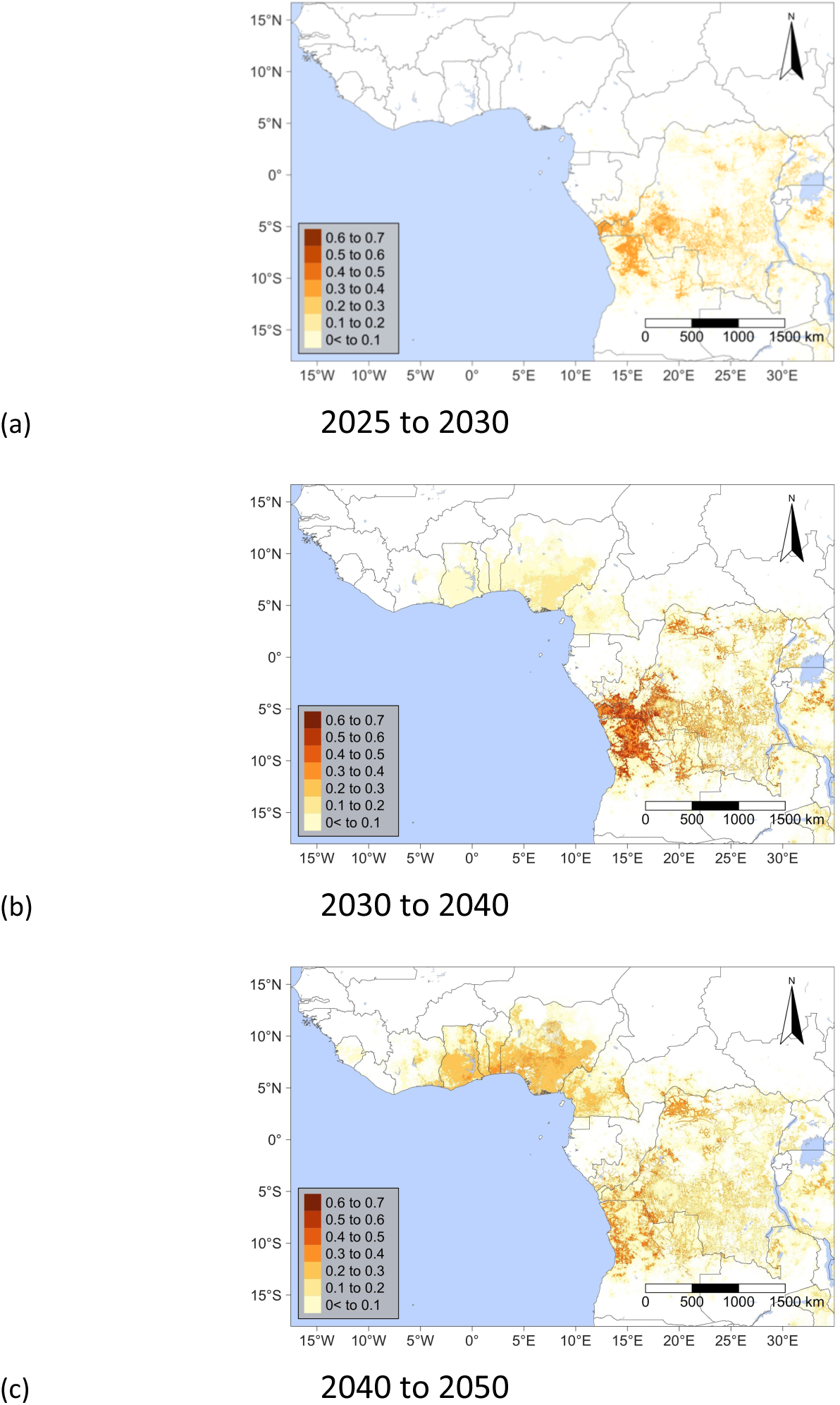
Differential risk maps representing the change in risk probability across the risk maps in Figure 4, highlighting regions that are likely to experience the most significant increase in the probability of CBSD occurrence across each time period.

### 2.4 Analysing the variability of predicted spread rates

The predicted arrival times of CBSI in each of thirty-two major cassava growing countries in SSA are summarised in (Figure 6). The arrival times are based upon simulations initialised in the endemic regions that also correspond to the Uganda surveillance data from 2004 and reported spread beyond Uganda (see methods). The variability in predictions reflect the intrinsic stochasticity of the epidemic model and the uncertainty in the estimated parameters. There is minimal variability for CBSI arrival times in countries neighbouring Uganda, with the uncertainty generally increasing for more distant countries.

**Figure 6:**
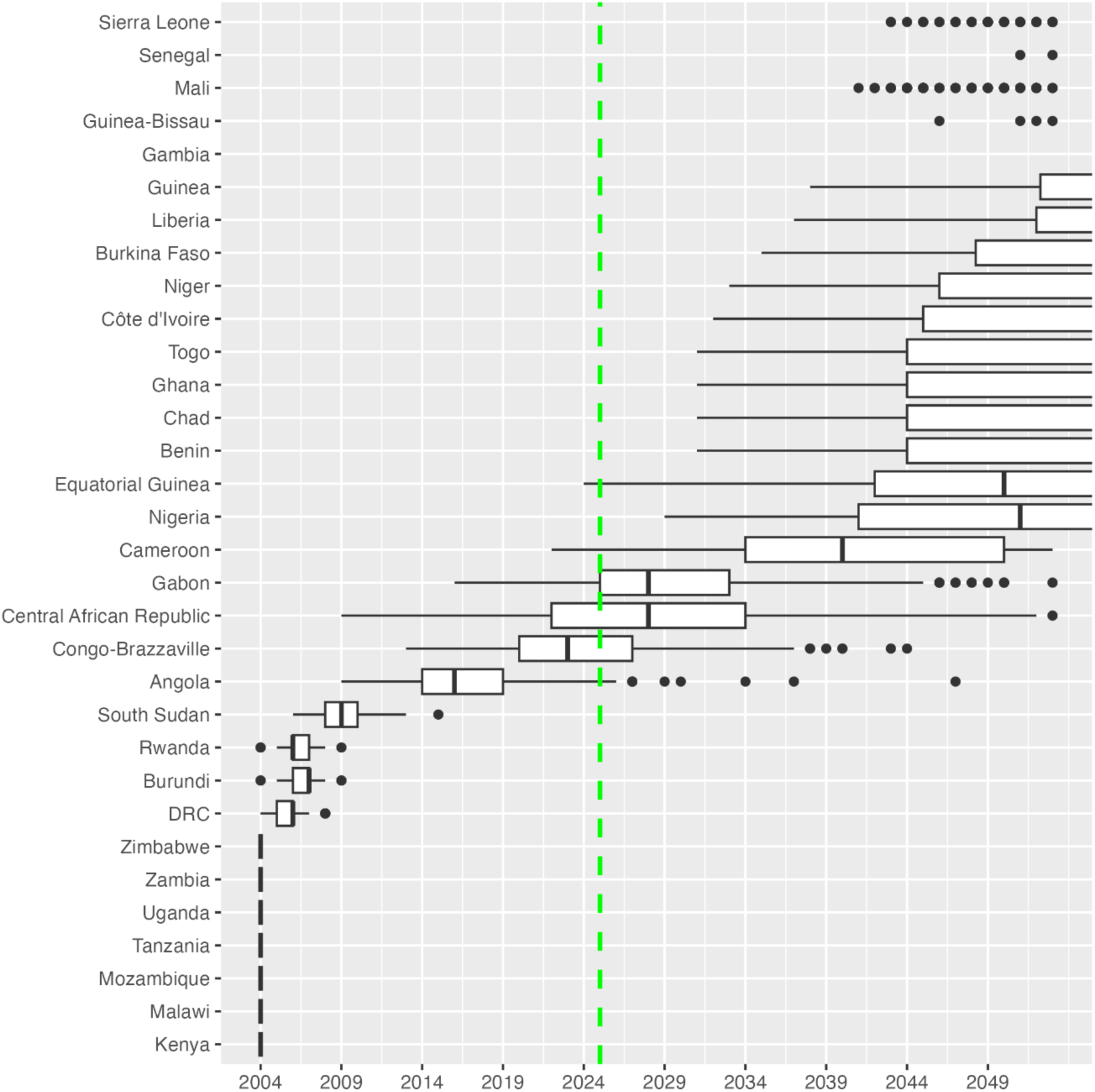
Box plot summarising the predicted year of CBSD arrival in the cassava growing countries of sub-Saharan Africa. The green line indicates the present day (2025). The central line corresponds to the median and the hinges correspond to the 25th and 75th percentiles. The whiskers extend to the largest and smallest values that fall within the 1.5 x IQR, where IQR is the inter-quartile range between the 25th and 75th percentiles. Points outside this range are plotted as outliers.

Simulations indicate a 45% probability of the epidemic arriving in Nigeria by 2050 (in the absence of any direct introduction event). There is a step change in the total size of epidemics that have reached Nigeria (Figure 7a), with Cameroon acting as a major gateway from Central into West Africa and Nigeria as a major booster in the rate and extent of epidemic spread. Supplementary Movie 2 presents simulations in a ranked order according to epidemic size in terms of the number of fields infected in 2050. The movie illustrates the relatively broad spatial variability in predicted epidemic spread amongst simulations, along with a representation of where the simulation values for two key dispersal parameters fall within the posterior distribution with predominantly smaller values for the dispersal parameters, α and ln(β), leading to more extensive westward spread. This trend in spread rates with dispersal parameters is highlighted in Figure 7b, showing the clustering of simulation parameters for which epidemics reached Nigeria at lower values of α.

**Figure 7:**
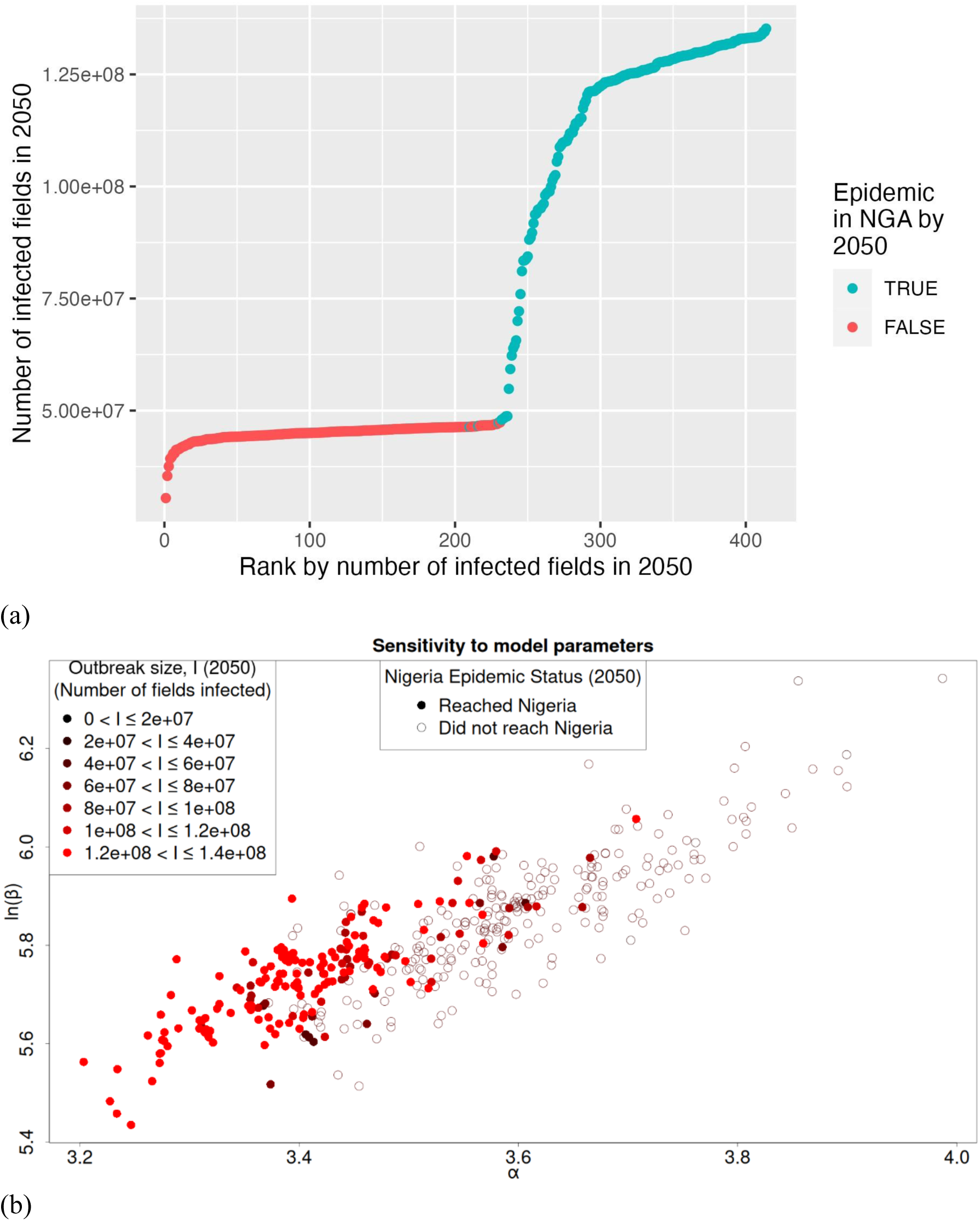
(a) The predicted size of the epidemic in 2050 in terms of the total number of fields in sub-Saharan Africa infected with CBSD. Each point represents a single simulation, coloured according to whether there are any CBSD infected fields in Nigeria by 2050. There are two distinct predicted scenarios in terms of the size of the resultant epidemic, with Cameroon acting as a major gateway from Central into West Africa and Nigeria as a major booster in the rate and extent of epidemic spread. (b) Effect of model parameters on predicted spread by 2050. The points represent samples of the (log) spread rate (ln(β)) and the dispersal kernel length scale (α) from the fitted posterior distribution. Filled circles indicate simulations in which the epidemic reaches Nigeria by 2050, and the shading of points indicates the severity of the outbreak, ranging from black for no outbreak to red being the most severe. The simulations in which the epidemic reaches Nigeria are predominantly associated with parameters with low α, corresponding to a longer scale for individual dispersal events.

## 3 Discussion

Since 2004, CBSD has spread from Uganda over 1000 km to Zambia in the south and to central DRC in the west, causing significant loss of yield for smallholder farmers in the region (Legg *et al*., 2015b). Given the importance of the threat posed to food security by the westward spread of the disease towards countries reliant on cassava as a staple crop, we adapted and extended a stochastic, epidemiological model that had previously been validated for the spread of CBSI across Uganda, to predict the present extent and future spread of CBSD in all thirty-two major cassava growing countries in SSA. The predictive accuracy of the extended model was validated relative to real-world reports of CBSD spread in East, Central and Southern Africa up to the present day, with the model showing strong correspondence to the timing of real-world observations (Figure 1b). These data on real world observations of the disease beyond Uganda were then incorporated into the analysis to improve predictive accuracy and reduce the uncertainty in model predictions (Figure 1c).

The predicted cross-continental epidemic spread rates are the result of the relatively local (i.e. < 50 km) combined action of the two CBSI dispersal mechanisms: the insect vector, *B. tabaci* and human-mediated movement of infected cuttings. The epidemic spread observed in Uganda and regionally in East Africa is consistent with the combined action of these two, relatively local, dispersal mechanisms. However, due to the long-term viability and persistence of the virus in cuttings, there is a risk of much longer-range introduction of CBSI to disease-free regions via cutting movement. Termed direct introduction, these movements can occur by air, sea or long-range road transport. To illustrate the scale of this risk for West Africa, we applied the model to direct introduction scenarios wherein the equivalent of five infected fields were introduced near each of the three major ports in Nigeria, Côte d’Ivoire, and Cameroon. The temporal rate of CBSD spread within a given country does vary significantly due to several factors including the density of the susceptible host, the connectivity of the landscape, and the abundance of the vector population in the region, with infection initially spreading more slowly from direct introductions in Côte d’Ivoire and Cameroon than Nigeria (Figure 3). Despite the varying local cassava densities and *B. tabaci* abundance, these three scenarios all indicate that epidemic spread would be far more rapid than seen to date in East Africa and result in extensive infection across West Africa, irrespective of the original source of introduction (Figures 2 and 3). As such, it is of vital importance to prevent introductions to West Africa. Nonetheless, even in the absence of spread in West Africa, all simulations predict extremely extensive epidemic spread throughout East and Central Africa, impacting millions of smallholders.

While the risk of direct introductions may be low with phytosanitary restrictions at seaports and airports, the potential harm of direct introductions could be extremely severe if the disease were to establish and spread prior to detection and response. We note that there are documented cases of large scale cutting movements by air for humanitarian purposes (ICRC, 2011). The risk of unintended consequences of importation of disease along with planting material merits careful assessment.

The scarcity of reports of CBSD epidemic expansion in East Africa and especially in Central Africa means there is significant uncertainty about the present-day distribution of the pathogen and disease. Therefore, for predictions of the future spread of the epidemic, the model could not be trivially initialised on the likely present-day extent of infection (see Methods). To arrive at an estimation of the present-day epidemic in East, Central and Southern Africa, simulations were initiated from the start of the epidemic in Uganda in 2004 and analysed to select simulations that predicted CBSI infected fields in the same location and timing as the available real-world reports beyond Uganda. Given the sparsity of reporting, we could only be confident that CBSI was present at the time of the report, as opposed to assuming CBSI absence prior to the first report. Matching simulations with reported presence allowed us to discard simulations that spread too slowly but did not allow us to impose an upper bound on those spreading too fast. However, we also imposed a constraint whereby selected simulations were also required to conform to surveillance data in Uganda from 2004-2017, which effectively introduced both upper and lower bounds on spread rates. By contrasting Figure 1b with Figure 1c, we see the improved prediction accuracy for arrival times in Central DRC by incorporating additional information on real-world observations.

Regarding the CBSD endemic region in coastal East Africa and Malawi, we assumed there was no fundamental distinction between the endemic and post-2004 epidemic regions, which is consistent with available data on the CBSI species distribution (Mbanzibwa *et al*., 2011; Ndunguru *et al*., 2015; Munganyinka *et al*., 2018; Casinga *et al*., 2018). Moreover, preliminary simulations with and without allowance for infection from the endemic coastal East African and Malawi regions indicated a limited contribution of the endemic region to the post-2004 epidemic spread. Nonetheless, to gain a complete picture of the spatial extent of CBSI infection in SSA, we included the endemic regions as sources of infection. However, data for the true distribution of infected fields in the endemic region as with information on the post-2004 epidemic spread were limited, which meant it was also necessary to make several assumptions to arrive at a credible distribution of infected fields in the endemic region.

In the absence of contrary evidence, we assumed that there is nothing fundamentally distinct about cassava cultivars or *B. tabaci* diversity in West Africa that would prohibit the spread of CBSI. Studies on the susceptibility of West African cassava varieties have demonstrated susceptibility to CBSI (Elegba *et al*., 2020; Ano *et al*., 2021). While there is not yet sufficient data, with further studies to both characterise the response of cassava cultivars to CBSIs and also to map the spatial distribution of cassava cultivars, this information could be incorporated into the model to increase local prediction accuracy. Whilst there is diversity in the *B. tabaci* species across SSA (Chen *et al*., 2019), we assume that all *B. tabaci* are equivalently efficient vectors, with the modulation in vector-driven spread rates resulting from the abundance of the vector, not the specific species in the complex. To the authors’ knowledge, no studies have explored the question as to whether there are differential CBSI transmission efficiencies across the species of the *B. tabaci* complex.

With major uncertainties in climate change, demographics, and agricultural trends we do not know how cassava production will change in the future. As a result, the model currently assumes a fixed level of cassava production across time. Also, despite incorporating data from over 18,000 surveys, the *B. tabaci* abundance records are also sparse at the continental scale, especially in Central Africa. Future studies could attempt to incorporate the impact of changing host production over time associated with human population growth on the spread rates of CBSI across the continent and mitigate the challenge of limited vector data by modelling the *B. tabaci* abundance explicitly. Further work is also underway to improve the inverse distance weighting (IDW) interpolation used to generate the vector abundance layer used here (see Methods). This work will incorporate weather data linked with a whitefly population dynamics model to produce a dynamic, environmentally driven whitefly abundance prediction. Moreover, the model currently integrates the two dispersal processes (i.e. vector and human-mediated cutting movement) into a single dispersal kernel due to the absence of data to independently parameterise these two processes (Godding *et al*., 2023). Future studies should attempt to collect sufficient data on cutting trade movements to begin to disentangle these two mechanisms.

The most pressing questions facing countries in disease-free regions is how to prepare for the threat of CBSD introduction in a way that makes the most efficient use of limited resources. The model described in this study not only provides estimates for arrival times across SSA but lays the foundations for a continental-scale quantitative framework wherein both surveillance and management options can be explored and optimised. The results presented here highlight the need for strong phytosanitary restrictions to prevent the direct introduction of infected planting material to West Africa, effective surveillance for early detection of CBSI incursions and rapid responses at appropriate scales. The mechanistic structure of the underlying epidemiological model used in the current analyses is well suited to compare scenarios for early detection and response. Phytosanitary treatments, such as quarantine, roguing, culling and the deployment of virus-free planting material as well as insect vector control can be easily simulated within the modelling framework and the likely dynamical effectiveness of different scenarios compared.

## 4 Methods

### 4.1 Extending the data-driven layers of the CBSD model

We take a parameterised model of the CBSD epidemic in Uganda (Godding *et al*., 2023) and extend the data-driven input layers using additional data on cassava host production and *B. tabaci* abundance from across SSA. For the host landscape layer, we use the CassavaMap model (Szyniszewska, 2020), which covers the 32 major cassava producing countries in SSA and convert from production volume per km^2^ to number of fields per km^2^ (Szyniszewska, 2020; Godding *et al*., 2023). When referring to a cassava field, we use a representative unit field size of 0.1 ha, yielding approximately one tonne of cassava root material.

For the *B. tabaci* vector abundance data, we combine 18,340 vector abundance records from cassava disease surveys in 17 countries across SSA, reporting the average field-level abundance of *B. tabaci.* The vector abundance data were partially compiled from previous publications (Eni *et al*., 2018, 2020, 2021; Ateka *et al*., 2018; Chikoti *et al*., 2018; Cossa *et al*., 2018; Kanyange *et al*., 2018; Mbewe *et al*., 2018; Tairo *et al*., 2018; Zeyimo *et al*., 2019; Alicai *et al*., 2019; Muhindo *et al*., 2020b; Oppong *et al*., 2020; Soro *et al*., 2021; Houngue *et al*., 2022; Doungous *et al*., 2022) (see Table 1 for details), with additional unpublished data provided by co-authors. These data were collected using the same in-field protocol as the Ugandan surveillance data and the relative vector abundance layer was generated using inverse distance weighted (IDW) interpolation with a power value of 1.0 with a 5 km resolution at the same spatial extent as the host production layer (Godding *et al*., 2023).

**Table 1:** Summary of the number of field records used to generate the *B. tabaci* abundance model layer. These data were partially compiled from multiple publications along with previously unpublished data provided by co-authors. A subset of these data was also used to seed locations for CBSD in the endemic region of coastal East Africa and Malawi.

| Country | Number of field surveys | Date range of field surveys used in whitefly layer | Relevant citations |
| --- | --- | --- | --- |
| Benin | 232 | 2015-2022 | (Houngue *et al.*, 2022) |
| Burkina Faso | 276 | 2015-2022 | (Soro *et al.*, 2021) |
| Cameroon | 342 | 2020-2022 | (Doungous *et al.*, 2022) |
| Côte d'Ivoire | 853 | 2016-2022 |  |
| DRC | 1326 | 2009-2022 | (Zeyimo *et al.*, 2019; Muhindo *et al.*, 2020b) |
| Gabon | 227 | 2020-2023 | (Mouketou *et al.*, 2022) |
| Ghana | 701 | 2015-2022 | (Oppong *et al.*, 2020) |
| Kenya | 505 | 2009-2017 | (Ateka *et al.*, 2018) |
| Malawi | 296 | 2012-2015 | (Mbewe *et al.*, 2018) |
| Mozambique | 1226 | 2012-2017 | (Cossa *et al.*, 2018) |
| Nigeria | 2127 | 2015-2022 | (Eni *et al.*, 2018, 2020, 2021) |
| Rwanda | 348 | 2009-2017 | (Kanyange *et al.*, 2018) |
| Sierra Leone | 268 | 2020-2022 |  |
| Tanzania | 809 | 2009-2017 | (Tairo *et al.*, 2018) |
| Togo | 344 | 2015-2020 |  |
| Uganda | 7507 | 2004-2017 | (Alicai *et al.*, 2019) |
| Zambia | 953 | 2009-2019 | (Chikoti *et al.*, 2018) |

### 4.2 Initial conditions for the epidemic state in 2004

For all model simulations, we define *t* = 0 as 1st January 2004, which we take to be the start of the Ugandan CBSD epidemic. To generate a model layer representing the distribution of CBSD-infected fields across all cassava-producing countries in 2004, it was necessary to account for the pre-2004 CBSD endemic region. Cassava brown streak disease has been reported as endemic in coastal East Africa and the shores of Lake Malawi since the first scientific reports beginning in the 1930s (Story, 1936; Nichols, 1950; Hillocks & Jennings, 2003). For the coastal countries of Kenya, Tanzania and Mozambique, polygons were created covering a 2 decimal degree wide region from the coast (approximately 220 km). In the case of Malawi, due to its smaller size, the entire country polygon was used. These polygons were overlaid with the surveillance dataset (described in the previous section) to identify CBSD-positive fields observed within the endemic region (Supplementary Figure 1a).

To create a layer that represents the distribution of infected fields in the endemic region, we ran preliminary simulations to extend the number of infected fields beyond the observations of CBSD positive fields from the surveillance data. For the initial conditions in these preliminary simulations, the equivalent of a single infected field was seeded at the location of each of the CBSD positive surveys from the surveillance data. An ensemble of 100 preliminary simulations were run using these initial conditions for 20 years. Simulations that exceeded 1% of infected fields in any non-endemic country were discarded, along with simulations that did not exceed 50% of fields infected in each of the endemic polygons. The value of 50% was selected due to the proportion of CBSD positive field surveys in the endemic polygons being approximately 50%. From the subset of simulations that met this criterion, a simulation was selected at random, and the 20-year end state was taken as representative of the 2004 CBSD distribution in the endemic region (Supplementary Figure 1b).

Initial conditions for the 2004 state were derived by combining the endemic region infection layer with the initial CBSD observations in Uganda near Kampala in 2004 (Alicai *et al*., 2019).

### 4.3 Running simulations from 2004 to 2054

Taking the initial conditions representing the epidemic state in 2004, we ran an ensemble of 10,000 simulations for a 50-year period using the University of Cambridge HPC service. Due to the high computational resource requirements of these simulations, we applied a one-month maximum time to completion cut off. Of the 10,000 simulations, 9,072 completed within this time limit.

### 4.4 Initial conditions for predictions beyond the present day

We use the observations of CBSD spread within and beyond Uganda to isolate the subset of stochastic simulations that correspond to the real-world observations up to the present day (Figure 1a). In doing so, we constrain the predictions of epidemic spread beyond the present day to be consistent with available information about the state of the system up to the present day. We take this approach as we do not know the true initial conditions for the model in the present day due to the sparsity of real-world observations and to address the challenge of extrapolating from surveillance to the true underlying state.

We utilise the simulated surveillance features of the model to simulate CBSD surveillance at the same time and locations as the real-world Ugandan surveillance (Alicai *et al*., 2019). We then calculate the two summary statistics, *S_kam_* and *S_nat_* covering the full dataset period of 2004-2017, referred to collectively as *S_inf_*, and isolate simulations that satisfy a tolerance of ε*_inf_* (Godding *et al*., 2023).

For the set of observations beyond Uganda (Figure 1a), we identify simulations that have any CBSD infected fields within a polygon corresponding to the area of the real-world CBSD observations by the end of the year of real-world observation. For the two reports from single sites with a specified latitude-longitude coordinate, labelled DRC (East) and DRC (Central) in Figure 1a, circular polygons were created with a radius of 100 km around the specified coordinate; for Rwanda and Burundi, the entire country polygons are used; for DRC (southeast), a polygon of Pweto territory was used; and for Zambia, we created a single polygon from the union of Chiengi and Kaputa districts.

Analyses are then performed on the subset of simulations that meet all specified requirements (i.e. correspond to Ugandan surveillance data and reported spread beyond Uganda). Out of the 9,072 simulations, a total of 414 met all these criteria.

Simulations of direct introductions to West Africa were chosen to have five independent infected field sites, as this was the size of the outbreak discovered in Uganda in 2004.

### 4.5 Code availability

The code is available at https://github.com/dsg38/cbsd_scenarios/

### 4.6 Data availability

The field record datasets collected in East Africa as part of the Cassava Diagnostics Project (CDP) are made available as part of the code repository. The remaining field datasets provided by the Central and West Africa by the Central and West African Virus Epidemiology Center (WAVE) are available upon reasonable request from WAVE.

## Acknowledgements

The authors gratefully acknowledge financial support from the Gates Foundation (Grant Number: INV-070408), the UK Foreign, Commonwealth and Development Office and the UK Biotechnology and Biology Research Council (BBSRC). We also acknowledge many helpful discussions and support from members of the Epidemiology & Modelling Group in Cambridge, in addition to partners from the Cassava Diagnostics Project (CDP) and the Central and West African Virus Epidemiology (WAVE) project.

This work was performed using resources provided by the Cambridge Service for Data Driven Discovery (CSD3) operated by the University of Cambridge Research Computing Service (www.csd3.cam.ac.uk), provided by Dell EMC and Intel using Tier-2 funding from the Engineering and Physical Sciences Research Council (capital grant EP/T022159/1), and DiRAC funding from the Science and Technology Facilities Council (www.dirac.ac.uk).

## Author contributions

CAG conceived the original project. DSG, ROJHS and CAG planned the modelling approach in response to discussions with JSP and AOE. DSG and ROJHS developed, tested and ran the code for the model simulations. DSG, ROJHS and CAG drafted the paper. All remaining co-authors conducted cassava disease surveillance, contributing datasets for the generation of the vector abundance layer and CBSD initial conditions, and reviewed the paper.

## Competing interests

The authors declare no competing financial interests.

## Notes

### Competing Interest Statement

The authors have declared no competing interest.

## References

Alicai T, Omongo CA, Maruthi MN et al., 2007. Re-emergence of cassava brown streak disease in Uganda. Plant Disease 91, 24–29.

Alicai T, Szyniszewska AM, Omongo CA et al., 2019. Expansion of the cassava brown streak pandemic in Uganda revealed by annual field survey data for 2004 to 2017. Scientific Data 6, 1–8.

Ano CU, Ochwo-Ssemakula M, Ibanda A et al., 2021. Cassava brown streak disease response and association with agronomic traits in elite Nigerian cassava cultivars. Frontiers in Plant Science 12.

Ateka EM, Onguso J, Mwaura SK et al., 2018. The cassava diagnostics project: a review of 10 years of research, Kenya. In: *The Cassava Diagnostics Project*: *A Review of 10 Years of Research*. 115–156.

Casinga CM, Monde G, Shirima RR, Legg J, 2018. First report of mixed infection of cassava brown streak virus and Ugandan cassava brown streak virus on cassava in north-eastern Democratic Republic of Congo. Plant Disease.

Casinga CM, Shirima RR, Mahungu NM et al., 2020. Expansion of the cassava brown streak disease epidemic in eastern Democratic Republic of Congo. Plant Disease.

Chen W, Wosula EN, Hasegawa DK et al., 2019. Genome of the African cassava whitefly *Bemisia tabaci* and distribution and genetic diversity of cassava-colonizing whiteflies in Africa. Insect Biochemistry and Molecular Biology 110, 112–120.

Chikoti P, Tembo M, Mirriam C et al., 2018. The cassava diagnostics project: a review of 10 years of research, Zambia. In: *The Cassava Diagnostics Project*: *A Review of 10 Years of Research*. 245–280.

Cossa N, Marcia R, Surge A et al., 2018. The cassava diagnostics project: a review of 10 years of research, Mozambique. In: *The Cassava Diagnostics Project*: *A Review of 10 Years of Research*. 181–204.

Day R, Abrahams P, Bateman M et al., 2017. Fall armyworm: impacts and implications for Africa. Outlooks on Pest Management 28, 196–201.

Doungous O, Masky B, Levai DL et al., 2022. Cassava mosaic disease and its whitefly vector in Cameroon: Incidence, severity and whitefly numbers from field surveys. Crop Protection 158, 106017.

Elegba W, Gruissem W, Vanderschuren H, 2020. Screening for resistance in farmer-preferred cassava cultivars from Ghana to a mixed infection of CBSV and UCBSV. Plants 9, 1026.

Eni AO, Efekemo OP, Onile-ere OA, Pita JS, 2020. South West and North Central Nigeria: Assessment of cassava mosaic disease and field status of African cassava mosaic virus and East African cassava mosaic virus. Annals of Applied Biology **n/a**.

Eni AO, Efekemo OP, Onile-ere OA, Pita JS, 2021. Survey dataset on the epidemiological assessment of cassava mosaic disease in South West and North Central regions of Nigeria reveals predominance of single viral infection. Data in Brief 38, 107282.

Eni AO, Efekemo OP, Soluade MG, Popoola SI, Atayero AA, 2018. Incidence of cassava mosaic disease and associated whitefly vectors in South West and North Central Nigeria: Data exploration. Data in Brief 19, 370–392.

FAO, 2021. Desert locust upsurge: progress report on the response in the Greater Horn of Africa and Yemen, January-April 2021. Rome, Italy: FAO.

FAO, 2022. FAOSTAT Statistics Database.

Godding D, Stutt ROJH, Alicai T, Abidrabo P, Okao-Okuja G, Gilligan CA, 2023. Developing a predictive model for an emerging epidemic on cassava in sub-Saharan Africa. Scientific Reports 13, 12603.

Goergen G, Kumar PL, Sankung SB, Togola A, Tamò M, 2016. First report of outbreaks of the fall armyworm *Spodoptera frugiperda* (J. E Smith) (Lepidoptera, Noctuidae), a new alien invasive pest in West and Central Africa. PLOS ONE 11, e0165632.

Gomez y, Paloma S, Riesgo L, Louhichi K (Eds.), 2020. The Role of Smallholder Farms in Food and Nutrition Security. Cham: Springer International Publishing.

Harvey CA, Rakotobe ZL, Rao NS et al., 2014. Extreme vulnerability of smallholder farmers to agricultural risks and climate change in Madagascar. Philosophical Transactions of the Royal Society B: Biological Sciences 369, 20130089.

Hillocks R, Jennings D, 2003. Cassava brown streak disease: A review of present knowledge and research needs. International Journal of Pest Management 49, 225–234.

Houngue JA, Houédjissin SS, Ahanhanzo C, Pita JS, Houndénoukon MSE, Zandjanakou-Tachin M, 2022. Cassava mosaic disease (CMD) in Benin: Incidence, severity and its whitefly abundance from field surveys in 2020. Crop Protection 158, 106007.

ICRC, 2011. ICRC Audiovisual Archives Picture/110534. ICRC Audiovisual archives.

Kanyange MC, Gashaka G, Munganyinka E et al., 2018. The cassava diagnostics project: a review of 10 years of research, rwanda. In: *The Cassava Diagnostics Project*: *A Review of 10 Years of Research*. 157–180.

Legg JP, Jeremiah SC, Obiero HM et al., 2011. Comparing the regional epidemiology of the cassava mosaic and cassava brown streak virus pandemics in Africa. Virus Research 159, 161–170.

Legg JP, Kumar PL, Kanju EE, Tennant P, Fermin G, Others, 2015a. Cassava brown streak. Virus Diseases of Tropical and Subtropical Crops, 42.

Legg JP, Lava Kumar P, Makeshkumar T et al., 2015b. Cassava virus diseases: biology, epidemiology, and management. Advances in Virus Research 91, 85–142.

Mbanzibwa DR, Tian YP, Tugume a. K et al., 2011. Simultaneous virus-specific detection of the two cassava brown streak-associated viruses by RT-PCR reveals wide distribution in East Africa, mixed infections, and infections in Manihot glaziovii. Journal of Virological Methods 171, 394–400.

Mbewe W, Mhone A, Semu R et al., 2018. The cassava diagnostics project: a review of 10 years of research, Malawi. In: *The Cassava Diagnostics Project*: *A Review of 10 Years of Research*. 205–204.

Meynard CN, Gay P-E, Lecoq M, Foucart A, Piou C, Chapuis M-P, 2017. Climate-driven geographic distribution of the desert locust during recession periods: subspecies’ niche differentiation and relative risks under scenarios of climate change. Global Change Biology 23, 4739–4749.

Mouketou A, Koumba AA, Gnacadja C, Zinga-Koumba CR, Abessolo Meye C, Ovono APM, Sevidzem SL, Mintsa R, Lepengué AN, Mavoungou JF, 2022. Cassava mosaic disease incidence and severity and whitefly vector distribution in Gabon. African Crop Science Journal, 30(2): 167–183.

Muhindo H, Wembonyama F, Yengele O et al., 2020a. Optimum time for harvesting cassava tubers to reduce losses due to cassava brown streak disease in Northeastern DRC. Journal of Agricultural Science 12, 70.

Muhindo H, Yasenge S, Casinga C et al., 2020b. Incidence, severity and distribution of cassava brown streak disease in northeastern Democratic Republic of Congo (F Yildiz, Ed,). Cogent Food & Agriculture 6, 1789422.

Mulenga RM, Boykin LM, Chikoti PC, Sichilima S, Ng’uni D, Alabi OJ, 2018. Cassava brown streak disease and Ugandan cassava brown streak virus reported for the first time in Zambia. Plant Disease 102, 1410–1418.

Mulimbi W, Phemba X, Assumani B et al., 2012. First report of Ugandan cassava brown streak virus on cassava in Democratic Republic of Congo. New Disease Reports 26, 11–11.

Munganyinka E, Ateka EM, Kihurani AW et al., 2018. Cassava brown streak disease in Rwanda, the associated viruses and disease phenotypes. Plant Pathology 67, 377–387.

Mware BO, Ateka EM, Songa JM, 2009. Transmission and distribution of cassava brown streak virus disease in cassava growing areas of Kenya. Journal of Applied Biosciences, 864–870.

Ndunguru J, Sseruwagi P, Tairo F et al., 2015. Analyses of twelve new whole genome sequences of cassava brown streak viruses and Ugandan cassava brown streak viruses from East Africa: diversity, supercomputing and evidence for further speciation. PLoS One 10, e0139321.

Nichols RFW, 1950. The brown streak disease of cassava. The East African Agricultural Journal 15, 154–160.

Omongo CA, Opio SM, Bayiyana I et al., 2022. African cassava whitefly and viral disease management through timed application of imidacloprid. Crop Protection 158, 106015.

Oppong A, Prempeh RNA, Abrokwah LA et al., 2020. Cassava Mosaic Virus Disease in Ghana: Distribution and Spread. In Review.

Retkute R, Hinton RGK, Cressman K, Gilligan CA, 2021. Regional differences in control operations during the 2019-2021 desert locust upsurge. Agronomy 11, 2529.

Sheat S, Fuerholzner B, Stein B, Winter S, 2019. Resistance against cassava brown streak viruses from Africa in cassava germplasm from South America. Frontiers in Plant Science 10, 567.

Soro M, Tiendrébéogo F, Pita JS et al., 2021. Epidemiological assessment of cassava mosaic disease in Burkina Faso. Plant Pathology 70, 2207–2216.

Story H, 1936. Virus diseases of East African plants. VI-A progress report on studies of the disease of cassava. East African Agricultural Journal 2, 34--39.

Szyniszewska AM, 2020. CassavaMap, a fine-resolution disaggregation of cassava production and harvested area in Africa in 2014. Scientific Data 7, 1–5.

Szyniszewska AM, Chikoti PC, Tembo M, Mulenga R, van den Bosch F, McQuaid CF, 2019. Cassava planting material movement and grower behaviour in Zambia: implications for disease management. bioRxiv, 528851.

Tairo F, Sseruwagi P, Kayuki C et al., 2018. The cassava diagnostics project: a review of 10 years of research, Tanzania. In: *The Cassava Diagnostics Project*: *A Review of 10 Years of Research*. 15–79.

Tomlinson, K., Bailey, A., Alicai, T., Seal, S., Foster, G., 2017. Cassava brown streak disease: historical timeline, current knowledge and future prospects. Molecular Plant Pathology 19, 1282–1294.

Tumwegamire S, Kanju E, Legg J et al., 2018. Exchanging and managing in-vitro elite germplasm to combat Cassava Brown Streak Disease (CBSD) and Cassava Mosaic Disease (CMD) in Eastern and Southern Africa. Food Security 10, 351–368.

Zeyimo B, Pita J, Kuhima M et al., 2019. Assessing the severity and the incidence of cassava root necrosis disease (CNRD) in western Democratic Republic of Congo. *International Journal of Agriculture*, Environment and Bioresearch 04, 237–253.

