## Supplementary material for "Predicting the cross-continental spread of the cassava brown streak disease epidemic in sub-Saharan Africa": S1 Supplementary Methods

### Endemic regions pre-2004

Regions with endemic infection of CBSI prior to the post-2004 CBSD epidemic in Uganda are recorded in Figure S1.

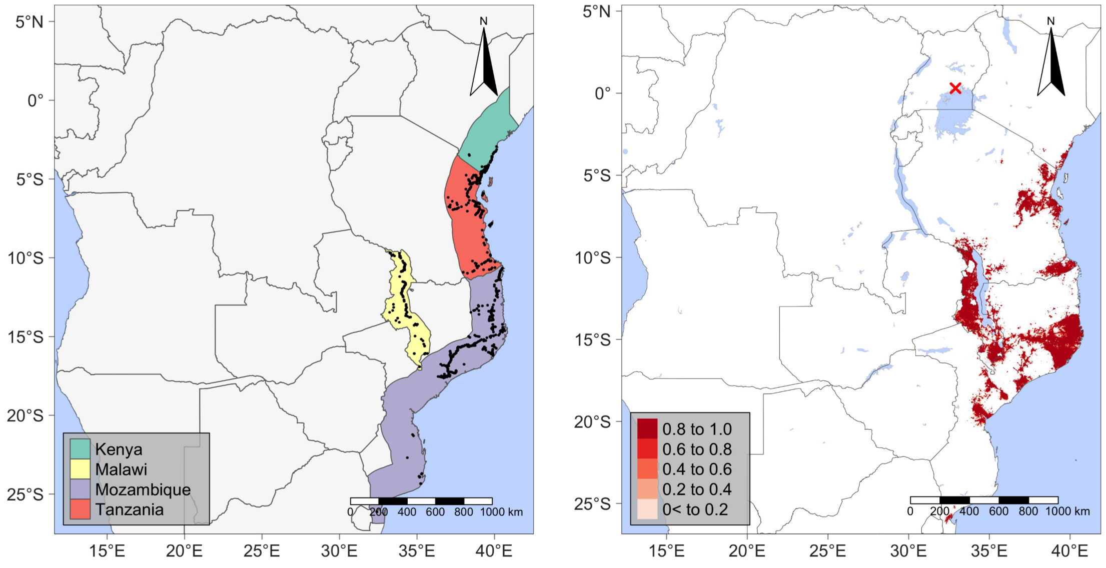


(a) (b)

Supplementary Figure 1: (a): An overview of the regions that are recorded as endemic for CBSD prior to the post-2004 CBSD epidemic in Uganda. Black points represent the locations of CBSD-positive field surveys used to seed infection to generate a realistic layer of infection for the 2004 state of CBSD infection in the endemic region. (b): The initial conditions taken to represent the distribution of CBSD infected fields at the start of the 2004 Ugandan CBSD epidemic. Red gradient indicates the proportion of cassava fields in a given 1 $km^2$ raster cell that are infected with CBSD. Red cross indicates the site of the initial 6 fields recorded as CBSD positive in Uganda.
